# Perigenual and Subgenual Anterior Cingulate Afferents Converge on Common Pyramidal Cells in Amygdala Subregions of the Macaque

**DOI:** 10.1101/2021.05.07.443119

**Authors:** EA Kelly, VK Thomas, A Indraghanty, JL Fudge

**Affiliations:** Department of Neuroscience, University of Rochester, Rochester NY 14642; Department of Pharmacology and Physiology, University of Rochester, Rochester NY 14642; Department of Psychiatry, University of Rochester, Rochester NY 14642

## Abstract

The subgenual (sgACC) and pregenual (pgACC) anterior cingulate are important afferents of the amygdala, with different cytoarchitecture, connectivity, and function. The sgACC is associated with arousal mechanisms linked to salient cues, while the pgACC is engaged in conflict decision-making, including in social contexts. After placing same-size, small volume tracer injections into sgACC and pgACC of the same hemisphere in male Macaques, we examined anterogradely labeled fiber distribution to understand how these different functional systems communicate in the main amygdala nuclei at both mesocopic and cellular levels. The sgACC has broad-based termination patterns. In contrast, the pgACC has a more restricted pattern which was always nested in sgACC terminals. Terminal overlap occurred in subregions of the accessory basal and basal nuclei, which we termed ‘hotspots’. In triple-labeling confocal studies, the majority of randomly selected CAMKIIα (+) cells (putative amygdala glutamatergic neurons) in ‘hotspots’ received dual contacts from the sgACC and pgACC. The ratio of dual contacts occurred over a surprisingly narrow range, suggesting a consistent, tight balance of afferent contacts on postsynaptic neurons. Large boutons, which are associated with greater synaptic strength, were approximately 3 times more frequent on sgACC versus pgACC axon terminals in ‘hotspots’, consistent with a fast ‘driver’ function. Together, the results reveal a nested interaction in which pgACC (’conflict/social monitoring’) terminals converge with the broader sgACC (’salience’) terminals at both the mesoscopic and cellular level. The pre-synaptic organization in ‘hotspots’ suggest that shifts in arousal states can rapidly, and flexibly influence decision-making functions in the amygdala.

**Significance statement:** The subgenual (sgACC) and perigenual cingulate (pgACC) have distinct structural and functional characteristics and are important afferent modulators of the amygdala. The sgACC is critical for arousal, while the pgACC mediates conflict-monitoring, including in social contexts. Using dual tracer injections in the same monkey, we found that sgACC inputs broadly project in the main amygdala nuclei, whereas pgACC inputs were more restricted and nested in zones containing sgACC terminals (‘hotspots’). The majority of CAMKIIα + (excitatory) amygdala neurons in ‘hotspots’ received converging contacts, which were tightly balanced. pgACC and sgACC afferent streams are therefore highly interdependent in these specific amygdala subregions, permitting ‘internal arousal’ states to rapidly shape responses of amygdala neurons involved in conflict and social monitoring networks.

## Introduction

Emotional processing involves coding sensory data as biologically relevant, or ‘salient’, for the organism to survive. The amygdala codes the emotional relevance of complex sensory inputs through an intricate network of intrinsic and extrinsic connections. In humans and monkeys, the amygdala is especially sensitive to salient stimuli of a ‘social’ nature (i.e. facial expression, vocal expressions; Gothard et al., 2007; Rutishauser et al., 2011; Mosher et al., 2014; Wang et al., 2014; Rutishauser et al., 2015; Wang et al., 2017), which supports survival in highly interdependent social groups. The same neural ensembles in monkey amygdala can code reward associated with both nonsocial and social stimuli (Munuera et al., 2018), indicating a common or overlapping circuitry available for coding ‘salience’ and ‘social’ cue evaluation.

In both human and nonhuman primates, the amygdala is a heterogenous structure with evolutionary expansion of ‘cortical-like’ nuclei that communicate directly with the cortex (in contrast to the ‘extended amygdala’ structures, which mainly regulate hypothalamus and brainstem; Stephan et al., 1987). The basal and accessory basal nuclei of the amygdala are the key sites of higher processing through strong, reciprocal connections with the prefrontal cortex (PFC), in particular the anterior cingulate (ACC) (Carmichael and Price, 1995; Ghashghaei et al., 2007; Cho et al., 2013; Sharma et al., 2020). These ‘cortical-like’ nuclei contain pyramidal projection neurons and interneuron populations similar to the cortex.

In the human and monkey, amygdala-ACC networks are implicated in flexibly and appropriately modulating reward learning (Zhang et al., 2013), threat and extinction learning (Klavir et al., 2013; Reddan et al., 2018) and learning from social cues (Allsop et al., 2018; Munuera et al., 2018; Dal Monte et al., 2020). The ACC-amygdala connection develops gradually over childhood and adolescence (Gabard-Durnam et al., 2014; Gee et al., 2018), is influenced by early life experience, and is vulnerable to dysregulation in a host of human psychiatric illnesses (Johansen-Berg et al., 2008; Kim et al., 2011; Burghy et al., 2012).

The primate ACC, including that in human, has multiple subdivisions based on cytoarchitectural, connectional, and functional features (Vogt et al., 2005). The ‘subgenual’ ACC (sgACC), and ‘pregenual’ ACC (pgACC) are most connected with the amygdala (Cho et al., 2013; Sharma et al., 2020). The sgACC (areas 25/14c) is involved in monitoring emotional arousal and autonomic states (Vogt et al., 2005; Rudebeck et al., 2014). In contrast, the pregenual ACC (areas 24/32) is involved in ‘conflict’ decision-making, particularly in social contexts (Modirrousta and Fellows, 2008; Amemori and Graybiel, 2012; Livneh et al., 2012; Apps et al., 2016; Lockwood and Wittmann, 2018; Palomero-Gallagher and Zilles, 2018).

In a previous broad-based study that evaluated ‘top-down’ projections from the PFC to amygdala using retrograde tracer injections, we found that cortical inputs to the amygdala co-project with one another in a hierarchical manner, dictated by the relative granularity of the cortical region of origin (Cho et al., 2013). The least differentiated cortices (including the sgACC) form a ‘foundational circuit’ throughout the basal and accessory basal nuclei, upon which increasingly complex information from more differentiated cortical regions (e.g. pgACC) is ‘layered’ in more restricted amygdala subregions.

In the present study, we took an anterograde tracing approach to examine the idea that ‘social and conflict monitoring’ nodes of the ACC (i.e. the pgACC) that project to the amygdala would always occur in the context of more foundational ‘salience’ inputs (sgACC) in the amygdala. We placed injections into area 25/14c (sgACC ‘salience/arousal’) and area 24 and 32 (pgACC, ‘social and conflict monitoring’) nodes of the ACC in the same animal and examined their patterns of segregation and overlap in specific amygdala nuclei. Then, in regions where sgACC and pgACC terminals overlapped, we further investigated the patterns of convergence or segregation onto pyramidal cell populations using high resolution confocal techniques. Finally, we characterized terminal bouton size from each afferent source.

## Materials and Methods

### Experimental Design

A total of 7 injections sites in 5 animals were analyzed for this study: 2 single injections into the sgACC, 1 single injection into the pgACC, and 2 pairs of combined injections in the sgACC/pgACC (Table 1). Using the criteria of Carmichael and Price (Carmichael and Price, 1994), we designated injections that involved areas 25/14c as subgenual anterior cingulate (sgACC) and injections into area 32 and/or 24b as perigenual anterior cingulate (pgACC), respectively. A small injection of a different tracer was placed into the sgACC and pgACC of the same hemisphere in the macaque in two animals. After sectioning and processing of the brains for tracers using immunocytochemistry, we mapped the distribution of anterogradely labeled afferent fibers within amygdala subregions and used these macroscopic maps to identify regions of terminal segregation and overlap in the amygdala. We then conducted triple immunofluorescent analyses aimed at examining the relationship of axon contacts onto presumptive glutamatergic neurons in regions of terminal overlap. The relative size of synaptic boutons associated with the sgACC and pgACC in regions of overlap were conducted in single-labeled, adjacent sections using stereologic methods.

**Table 1.** Experimental cases.

| Case | Age<br>(yrs) | Wt<br>(kg) | Target | Hemisphere | Tracer | Tracer<br>Vol |
| --- | --- | --- | --- | --- | --- | --- |
| 37 | 3.7 | 4.3 | 25r | L | FR | 40 |
| **\*46** | 3.2 | 4.9 | 24b | L | FR | 40 |
| **\*46** | 3.2 | 4.9 | 25r/c | L | FS | 40 |
| 50 | 4.25 | 4.7 | 25r/c | L | FR | 40 |
| **\*53** | 5.9 | 5.5 | 32r/c | L | FS | 40 |
| **\*53** | 5.9 | 5.5 | 25c,14 | R | FR | 40 |
| 54 | 3.3 | 5 | 32r/c, 24b | R | FS | 150 |

\* denotes part of an injection pair

### Animals and surgery

Injections were stereotaxically placed in 5 male *Macaque fascicularis* (Worldwide Primates, Tallahassee, FL USA) weighing between 4.3kg and 5.5 kg (Table 1). Prior to surgery and tracer injection, a T2 anatomical MRI scan using a custom head coil (0.5 x 0.5 x 0.88 <u>millimeter</u> resolution) was acquired on each subject. Thus, each subject had unique coordinates based on individual anatomy. Seven days prior to surgery, animals began a daily course of perioperative gabapentin (25 mg/kg) for preventative pain management. On the day of surgery, the monkey was sedated with intramuscular ketamine (10mg/kg) and then intubated, maintained on 1.5% isoflurane, and stabilized in a surgical stereotaxic apparatus. A craniotomy was performed under sterile conditions. In four animals, small injections (40 nl) of bidirectional tracers tetramethylrhodamine (fluoruby, FR) and fluorescein (FS) were pressure injected over a ten-minute period into the sgACC and/or pgACC of the same hemisphere, using coordinates calculated from the T2 MRI atlas created for that animal. Because of the relatively small terminal field resulting from 40 nl injections in the pgACC (see Results), a single injection of 150 nl was pressure injected into pgACC of a fifth animal for comparison (case MF54FS). For all injections, the syringe was left in place for 20 minutes after each injection to prevent tracking of the tracer up the injection track. Only one tracer injection of each type was made per animal. Following placement of planned injections, the bone flap was replaced, and the surgical site was closed. Post-operative daily monitoring of animals for signs of discomfort was conducted and gabapentin tapered accordingly.

### Tissue preparation

Twelve to fourteen days post-surgery, the animals were placed into a deep coma with pentobarbital (initial dose 20 mg/kg via intravenous line). They were sacrificed by intra-cardiac perfusion using 0.9% saline containing 0.5 milliliters of heparin sulfate (200ml/min for 15-20 minutes), followed by cold 4% paraformaldehyde in 0.1M phosphate buffer, pH 7.2 (PB) (200ml/min for 15-20 minutes). Following brain extraction, brains were postfixed overnight in 4% paraformaldehyde solution, then submerged sequentially in 10%, 20%, and 30% sucrose solutions until they sank in each. Brains were sectioned on a freezing, sliding microtome into 40um sections. Each section was placed sequentially in 24-compartment slide boxes containing cold cryoprotectant solution (30% sucrose and 30% ethylene glycol in 0.1 M PBS) and stored at -20°C.

### Single label immunocytochemistry (ICC)

To assess the location of anterogradely labeled fibers in amygdala subregions, we first stained 1:8 sections through the amygdala for each tracer. We selected adjacent sections to stain with cresyl violet or immunostain for NeuN in order to localize injection sites in the frontal cortex. In the amygdala, adjacent sections were stained for acetylcholinesterase (AChE) activity. AChE histochemistry shows clear demarcations of nuclear boundaries in the non-human primate amygdala (Amaral and Bassett, 1989) (Fig. 1) For cases with dual injections, three 1:24 adjacent compartments of tissue through the amygdala were selected: one for the sgACC tracer, one for the pgACC tracer, and one intervening compartment for AChE staining. Sections selected for ICC were rinsed in 0.1MPB, pH 7.2, with 0.3% Triton-X (PB-TX) overnight. The following day, tissue was treated with endogenous peroxidase inhibitor for five minutes and then thoroughly rinsed with PB-TX, and placed in a blocking solution of 10% normal goat serum in PB-TX (NGS-PB-TX) for 30 minutes. Following rinses in PB-TX, tissue was incubated in primary antisera to FR (1:1000, Invitrogen, rabbit) or FS (1:2000, Invitrogen, rabbit) for ∼96 hours at 4°C. Tissues were then rinsed with PB-TX, blocked with 10% NGS-PB-TX, incubated in biotinylated secondary anti-rabbit antibody, and then incubated with avidin-biotin complex (Vectastain ABC kit, Vector Laboratories, Burlington, ON Canada). After rinsing, the tracer was visualized with 2,2’-diaminobenzidine (DAB, 0.05mg/ml in 0.1M Tris-buffer). Sections were mounted out of mounting solution (0.5% gelatin in 20% ETOH in double distilled water) onto subbed slides, dried for 3 days, and coverslipped with DPX mounting media (Electron Microscope Sciences, Hatfield, PA). Additional sections were also selected, immunostained and then counterstained with Nissl to confirm fiber localization in specific nuclei, including the intercalated cell islands. Immunostaining with NeuN was conducted similar to the procedure described above, using anti NeuN (1:5000, EMD Millipore, mouse) and the avidin biotin reaction.

**Figure 1.**
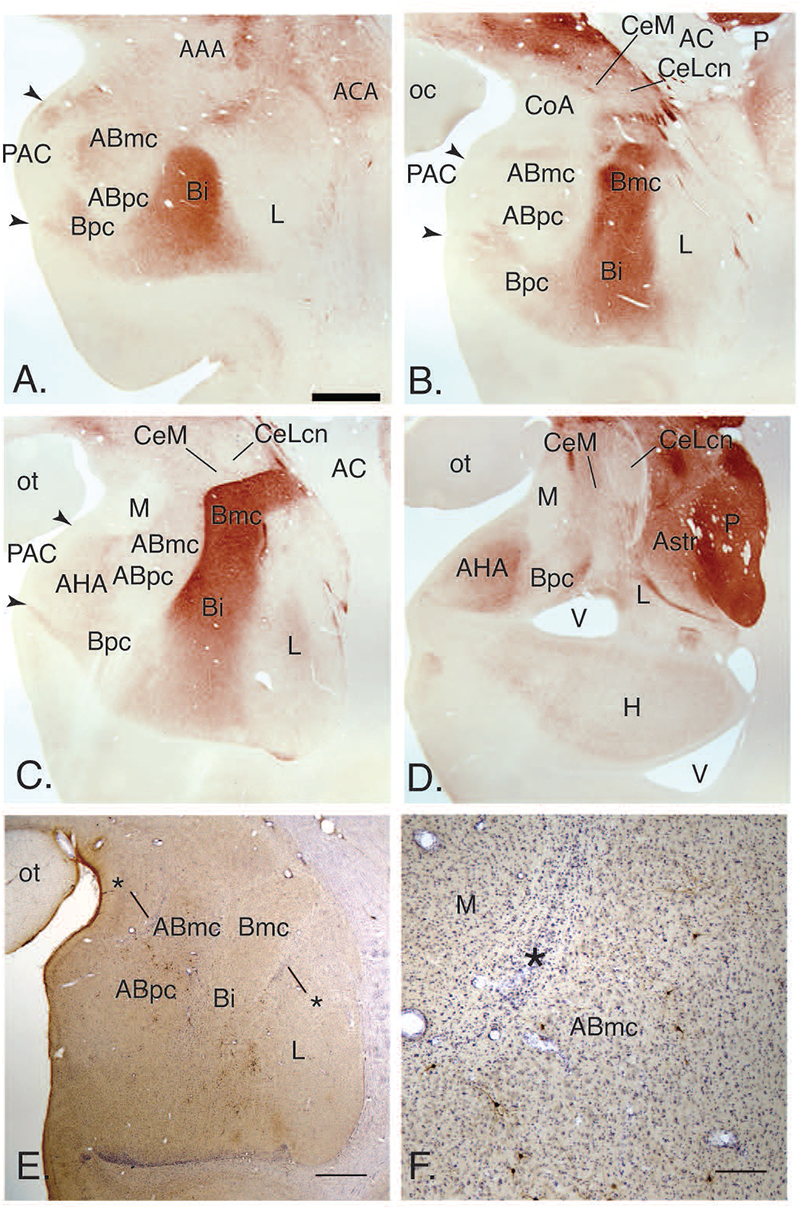
Histological markers for boundary designations. (A-D) Acetylcholinesterase (AChE) stained sections of *Macaque fasicularis* amygdala. (E-F) Identification of intercalated cell masses (ICMs) following anti-tracer immunohistochemistry with Nissl counterstain. * denote ICMs bordering amygdala nuclei. Abbreviations: **AAA**, anterior amygdaloid area; **ABmc**, magnocellular subdivision of the accessory basal nucleus; **ABpc**, parvocellular subdivision of the accessory basal nucleus; **AC**, anterior commissure; **AHA**, amygdalohippocampal area; **Astr**, amygdalostriatal area; **Bi**, intermediate division of the basal nucleus; **Bmc**, magnocellular division of the basal nucleus; **Bpc**, parvocellular division of the basal nucleus; **CeM**, central nucleus, medial subdivision; **CeLcn**, central nucleus, lateral subdivision; **CoA**, anterior cortical nucleus; **H**, hippocampus; **L**, lateral nucleus; **M**, medial nucleus; **oc**, optic chiasm; **ot**, optic tract; **P**, putamen; **PAC**, periamygdaloid complex; **V**, ventricle. Scale bar= A,E= 500 um; F= 200 um.

### Triple immuno-fluorescent labeling for tracers and CAMK-IIα

To determine the relationship of anterogradely labeled fibers from the peri- and subgenual ACC into the amygdala, and their relationship with glutamatergic amygdala neurons, we performed triple immunofluorescent staining for each tracer and calmodulin-dependent protein kinase II (CAMK-IIα)(McDonald et al., 2002) on an additional series through the amygdala in animals with paired injections. Optimization and specificity of fluorescent staining for all antigens was first conducted in single labeling experiments, with reference to the permanently labeled compartments for tracer (above) and CAMKIIα. Sections were rinsed in 0.1M PB, pH 7.2, with 0.3% Triton-X (PB-TX) overnight. The following day, tissue was treated with endogenous peroxidase inhibitor for 30 minutes and then thoroughly rinsed with PB-TX and placed in a blocking solution of 10% normal donkey serum in PB-TX (NDS-PB-TX) for 1 hour. Following rinses in PB-TX, tissue was incubated in 3% NDS-PB-TX primary antisera to FR (1:1000, Invitrogen #A6397, *made in rabbit*), FS (1:500, Invitrogen, #A11095, *made in goat*) and CAMK-IIα (1:1,000, Millipore #05-532, *made in mouse*) for ∼96 hours at 4°C. Tissues were then rinsed with PB-TX, blocked with 3% NDS-PB-TX and first incubated in biotinylated secondary anti-mouse antibody (1:200, Vector Labs, CAMK-IIα amplification) overnight at 4°C. Tissues were visualized following pooled incubation with donkey anti-rabbit Alexa Fluor 568 (1:200, FR visualization), donkey anti-goat Alexa Fluor 488 (1:100, FS visualization) and Streptavidin 647 (1:200, CAMK-IIα visualization). Tissue was mounted out of 0.1M PB, pH 7.2 and cover-slipped with Prolong Gold anti-fade mounting media (Invitrogen).

### Analysis

### Charting of anterograde fiber labeling in specific amygdala subregions

For all cases, charting of the distribution of anterogradely labeled fibers throughout the entire amygdala was first done on single-labeled, permanently stained tissue. Anterogradely labeled fibers for each case were visualized using an Olympus BX51 microscope equipped with a darkfield light source. Fibers were manually traced using an attached camera lucida drawing tube using a 4X and 10x objectives. Putative terminal fibers were characterized as thin processes with boutons. Thick labeled fibers without beaded processes were classified as fibers of passage and were not traced. Paper traces were scanned on a flat-bed scanner at high-resolution. Images were imported, stitched together and digitized using Adobe Photoshop CC. AChE-stained adjacent sections were projected onto anterograde traces using a JENA projector and nuclei borders were manually traced with the aid of landmarks within the tissue (i.e. blood vessels) and transferred onto digital files using a drawing tablet interfaced with the Adobe Illustrator CC. Tracer labeled sections and adjacent AChE labeled sections were placed into separate layers of in each file and aligned. Final image digitation and post-processing was performed using Adobe Illustrator CC. For paired injections in the same animal, the relationship of the anterogradely labeled fibers resulting from each injection site, and their localization in specific amygdala subregions, was done by turning off and on the visibility of layers. Regions of the main cortical-like nuclei that contained either segregation (‘non-overlap’) or overlap (‘hotspots’) of pgACC and sgACC labeled terminals were identified.

To quantify the density of anterograde fiber distribution in regions of overlap, we combined dark field microscopy with ImageJ (FIJI) analysis (Schneider et al., 2012). From cases with paired injections, low power images (4x objective) of ‘hotspots’ were first collected using dark field microscopy, and then re-captured at higher magnification (20X objective), using blood vessels and other landmarks to align ROIs across sections. Using Image J, the mean background density was determined from three unlabeled areas of each image. Images were manually thresholded to isolated fiber labeling (0-255 density scale; 0=black, 255=white), then converted to a binary scale (divided by 255; labeled regions=1, unlabeled regions=0). Original images were than multiplied by the binary masked image to obtain a density value for the image. The final density measurement was obtained after mean background subtraction.

#### Confocal capture and analysis of interactions between tracer-labeled fibers and CAMK-IIα positive cells

Cases with paired injections in the sgACC and pgACC and their regions of labeled fiber overlap in the amygdala (subregions that contained glutamatergic neurons) (‘hotspots’) were further assessed using high power (confocal) microscopy. We used a semi-automated algorithm to detect and quantify relative numbers of tracer labeled boutons on amygdala CAMK-IIα positive cells. While synaptic contacts can only be confirmed at the electron microscope level, we devised a method to examine the relationship of putative ‘contacts’ and their relationship to one another using confocal methods. Triple-immunofluorescent images were collected on a Nikon A1R HD Laser Scanning Confocal with NIS-Elements (Center for Advanced Light Microscopy and Nanoscopy) software using tissue landmarks on adjacent AChE-stained sections such as blood vessels for alignment, focusing on regions of interest (ROIs) where tracer labeled fibers overlapped at the light microscopic level. The following excitation lasers (ex) and emission filters (em) were used for confocal imaging: Alexa Fluor 488; ex 488, em 525/50, Alexa Fluor 568; ex 561, em 595/50, Streptavidin 647; ex 640, em 650 LP. Overview images were collected using a 20x/0.75 NA Nikon Plan Apochromat VC objective to locate ROIs in regions of tracer labeled fiber overlap, and z-series stacks were selected and collected using a 60x/ 1.49 NA Nikon Apochromat TIRF objective (xy pixel size of 0.17 micron; z-step size of 0.3 micron). 2-3 ROIs per ‘hotspot’ were collected per slide in each nucleus (n=2-3 slides; rostral to caudal extent of the amygdala) for each experimental case.

Projection images were analyzed with Imaris 9.6 software (Bitplane). Within each collected image stack (approximately 6-9 stacks per ROI per animal), ten CAMK-IIα positive cells were randomly selected for analysis. XYZ coordinate locations for each analyzed cell were carefully logged. In the 3D module, CAMK-IIα labeling was visualized using the “*surface rendering*” option (grain size=0.345 um diameter). The “smoothing” tool was disabled, as this introduces an artificial uniformity to the cell surface. Next, using the interactive software histogram of Cy5 voxels (volumetric pixels), a threshold was selected to include as much of the neuronal soma and proximal dendrites as possible while excluding any background. A second threshold (interactive size threshold) was next applied to exclude any further extraneous background labeling while still retaining the true volume of the labeled cell. Final renderings were then analyzed in 3D and looked at from all angles. In instances where the labeling was disjointed but clearly part of the actual cell, segments were linked, resulting in an n=10 cells per image.

Fibers were analyzed using the “*spot rendering*” option. In the “slice view”, fiber thickness was checked (a line measurement across the entire thickness of the fiber), resulting in a final size diameter of 2 um for all cases. The sensitivity for selected spots was adjusted using the automatically generated interactive histogram based on voxel size. We selected an area on the histogram to accurately detect as many boutons as possible without creating artifacts. Identical coordinates used during the “*surface rendering*” step were then applied to the “*spot rendering*” steps to ensure a 1:1 registration between the analyzed locations (CAMK-IIα containing cells) for detection and analysis of tracer labeled boutons. This step was conducted for each tracer labeled bouton, and individually overlaid on the CAMKIIα-positive surface rendering.

Once the combined CAMK-IIα ‘surface’ rendering and ‘spot’ renderings for tracer labeled boutons were constructed, we sought to determine the proportion of putative glutamatergic neurons contacted by tracer-labeled spots, including the proportion of CAMK-IIα containing neurons receiving dual-contacts. To do this, we used the “shortest distance” module to filter out all spots that were more than 0.5um from the cell surface. In the Imaris software, spot/surface contacts are based on a measurement from the center of the spot (radius) to the edge of the rendered surface object. As ‘spots’ were calculated as 2um, a radius of 1um would determine all spots “touching” a surface. Fiber boutons located close to CAMK-IIα cells but not in actual contact, may have an apparent voxel overlap due to the inherent resolution limits of light microscopy (a blurring of fluorescent edge resolution that extends beyond the actual true surface boundary). False-positive contacts can be controlled for by requiring a minimum number of overlapping voxels for an object to be classified as a true contact (Wouterlood et al., 2007). Our analysis was restricted to a maximum object to surface distance of 0.5um (at least a 50/50 overlap of spot and surface objects; Fig. 2) to create a relatively stringent inclusion criteria for assessing ‘synaptic contact’. Each CAMK-IIα (+) cell was analyzed in 3D and manual counts were performed in each fluorescent channel. We sought to determine a range of cell-contact types; including (i) no contacts, (ii) individual tracer contacts, and (iii) cells that contained dual contacts (single tracer contacts on the same cell). We present our results as (i) the proportion of pgACC/sgACC contacts on all CAMK-IIα cells and (ii) the percentage of contact type (see description above) onto CAMK-IIα (+) cells across amygdalar nuclei. For CAMK-IIα positive neurons receiving dual contact, we also analyzed the ratio of pgACC to sgACC contacts on each individual cell, presented as the percentage of neurons with different ratios of pgACC:sgACC contacts per ‘hotspot’ as well as the frequency of those contacts.

**Figure 2.**
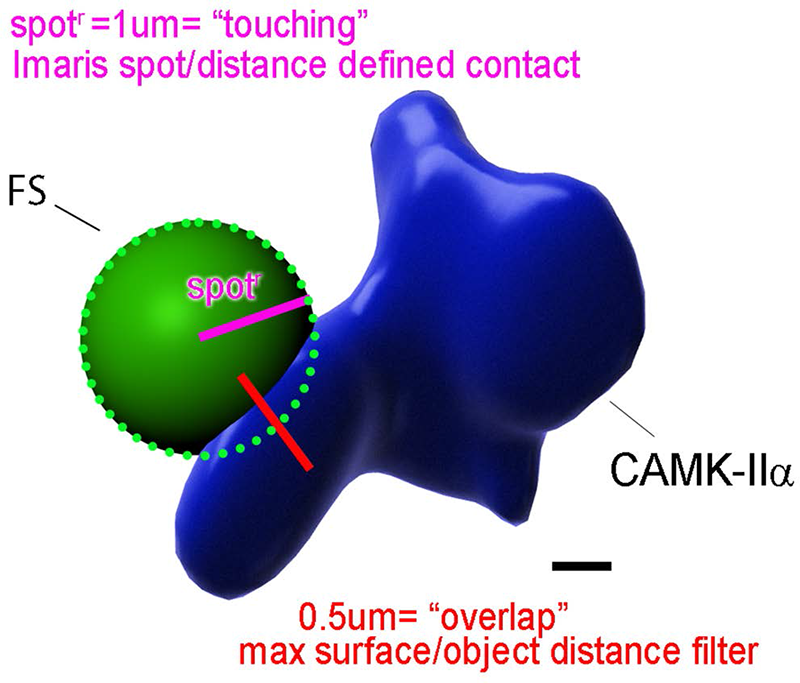
Triple immunofluorescence analysis: criteria for spot/surface contact. In Imaris software, spot/surface contacts are based on a measurement from the center of the spot (spot^r^) to the edge of the rendered surface object. As ‘spots” were calculated as 2um, a radius of 1um would determine all spots “touching” a surface (pink line). Our analysis was restricted to a maximum object to surface distance of 0.5um (red line) (approximately 50% overlap of spots and surfaces). Scale bar= 0.5 um.

#### Analysis of bouton size from pgACC and sgACC

To examine the relative size of axonal boutons from the pgACC and sgACC, we used unbiased stereologic methods to survey the basal and accessory basal nucleus ‘hotspots’ where overlapping terminals were found in each case. For comparison with ‘hotspots’, regions with non-overlapping sgACC (sgACC non-OL) terminals were also assessed (Fig. 5, dark blue). Using adjacent sections immunostained for the relevant tracer placed either in the sgACC or pgACC, the general terminal distribution in each ‘hotspot’ was drawn using a 2x (Plan, NA 0.05 ∞/-) objective. To sample under 100x (UPlanFl, NA 1.3) oil immersion objective, sampling parameters were: grid size 300 x 300, dissector height of 2um and a z-height of 5 um, resulting in sufficient sampling to yield a coefficient of error < 0.10. Axonal varicosities and terminal-like structures are considered synapses, as previous confirmed by electron microscopic methods (Ashaber et al., 2020). Both were counted using systematic, random sampling strategies in single-labeled adjacent slides for each tracer. The nucleator method (isotropic), which assumes sphericity of the structure, was applied to measure each putative synaptic structure along the Z axis, in the sampling frame (StereoInvestigator, Microbrightfield Bioscience, Williston, VT). Results were expressed in um^3^. Frequency histograms were created for each tracer in each area, and assessed for shape, spread, and percent of synaptic terminals > 0.52 μm^3^ (equivalent to 1 um diameter).

### Statistics

Statistical analyses were performed using Graphpad Prism software (V9.0.2 for Windows, LaJolla, California). A two-way ANOVA was used to compare tracer-labeled density measurement in regions of fiber overlap (‘hotspots”). A two-tailed unpaired student t-test was used to compare the total number of contacts in each injection group across amygdalar region. Similarly, a two-tailed unpaired student t-test was used to compare pgACC/sgACC ratio means across amygdalar nuclei. A 2x3 two-way ANOVA investigated the relationship of contact type on injection site (pgACC vs sgACC) and ROI (ABmc vs Bi). A two-way ANOVA was used to compare pgACC/sgACC ratio bin means across amygdala nuclei and injection sites. A one-way ANOVA was used to compare rostral/caudal differences in dual contacts. All statistical tests were corrected for multiple comparisons using Tukey’s multiple comparison test (Tukey’s HSD) where appropriate. P<0.5 was deemed statistically significant. Error bars are presented as standard error of the mean.

## Results

### Primate amygdala

The nonhuman primate amygdala spans 5-6 mm. in the rostrocaudal plane, and its nuclei shift in shape and cellular characteristics within this space (Fig. 1. A-D).

Acetylcholinesterase (AChE) is used to follow cytoarchitectural boundaries between and within nuclei (Amaral and Bassett, 1989). The ‘deep’ nuclear group of the amygdala includes the lateral, basal, and accessory basal nuclei, and also referred to as ‘cortical-like’ nuclei due to specific cellular and connectional characteristics (Carlsen and Heimer, 1988). These large nuclei contain pyramidal neurons and interneurons, and are intimately connected with broad regions of the prefrontal cortex (Ghashghaei et al., 2007; Cho et al., 2013). They are relatively expanded in primate species compared to the rodent, and with this expansion comes increased subnuclear organization which is only partially described here. The lateral nucleus is divided into dorsal, ventral intermediate, and ventral subdivisions with differing cellular features and AChE staining. The accessory basal nucleus consists of the magnocellular (ABmc) and parvicellular (ABpc) subdivisions, which have high and low levels of AChE, respectively. There is also a ‘sulcal’ subdivision of the accessory basal nucleus (ABs) that is medial and posterior relative to the other subdivisions. The basal nucleus is subdivided into magnocellular (Bmc), intermediate (Bi) and parvicellular subdivisions (Bpc), which have a decreasing gradient of cell size and AChE staining. The basal nucleus is surrounded by the paralaminar nucleus (PL), which is most prominent at its ventral boundary, but has almost absent AChE staining and not seen in these preparations. The intercalated cell islands are inhibitory neurons, that are lodged in the major fiber tracts that surround the ‘cortical-like’ nuclei. These small clusters of GABAergic neurons (Pitkanen and Amaral, 1993) also cannot be appreciated with AChE staining but are easily seen in cresyl violet preparations (Fig 1 E-F).

The anterior and posterior cortical nuclei (CoA and CoP, respectively), the medial nucleus, and the periamygdaloid cortex form the ‘superficial’ amygdala region. The amygdalohippocampal (AHA) region is an AChE-rich nucleus that forms a broad transitional region with the hippocampus, and is also composed of glutamatergic neurons and interneurons. In the dorsal amygdala, the central nucleus is situated caudally, and in is divided into medial and lateral central core subdivisions. It is separated from the striatum by a relatively broad, heterogeneous amygdalostriatal area (Fudge and Tucker, 2009). The medial and central amygdaloid nuclei are considered part of the ‘extended amygdala’ macrostructure, and are largely GABAergic.

### Injection site placement

The sgACC comprises areas 25c and 14c of the ACC; the pgACC comprises areas 32 and 24 (Fig. 3A and B, Table 1). The sgACC is agranular cortex that is defined by a lack of granular layer IV. sgACC injections were mainly in area 25, with slight involvement of 14c only in case 53FR. Case 37FR was located most rostrally, with Cases 50FR, 46FS, and 53FR at progressively caudal levels (Table 1). Injections into the pgACC, which is slightly more differentiated (Fig. 3A), were in either area 24b (Case 46FR) or area 32r/32c (Case 53FS) or encompassed both area 24b and the entire rostrocaudal extent of 32 in one case with a large single injection (Case 54FS). Figure 3C shows examples a single sgACC injection (case 37FR) and Figures 3-D1 shows dual pgACC injection (case 53FS) and sgACC injection (case 53FR), each with adjacent sections stained to localize placement with respect to cytoarchitectural profiles.

**Figure 3.**
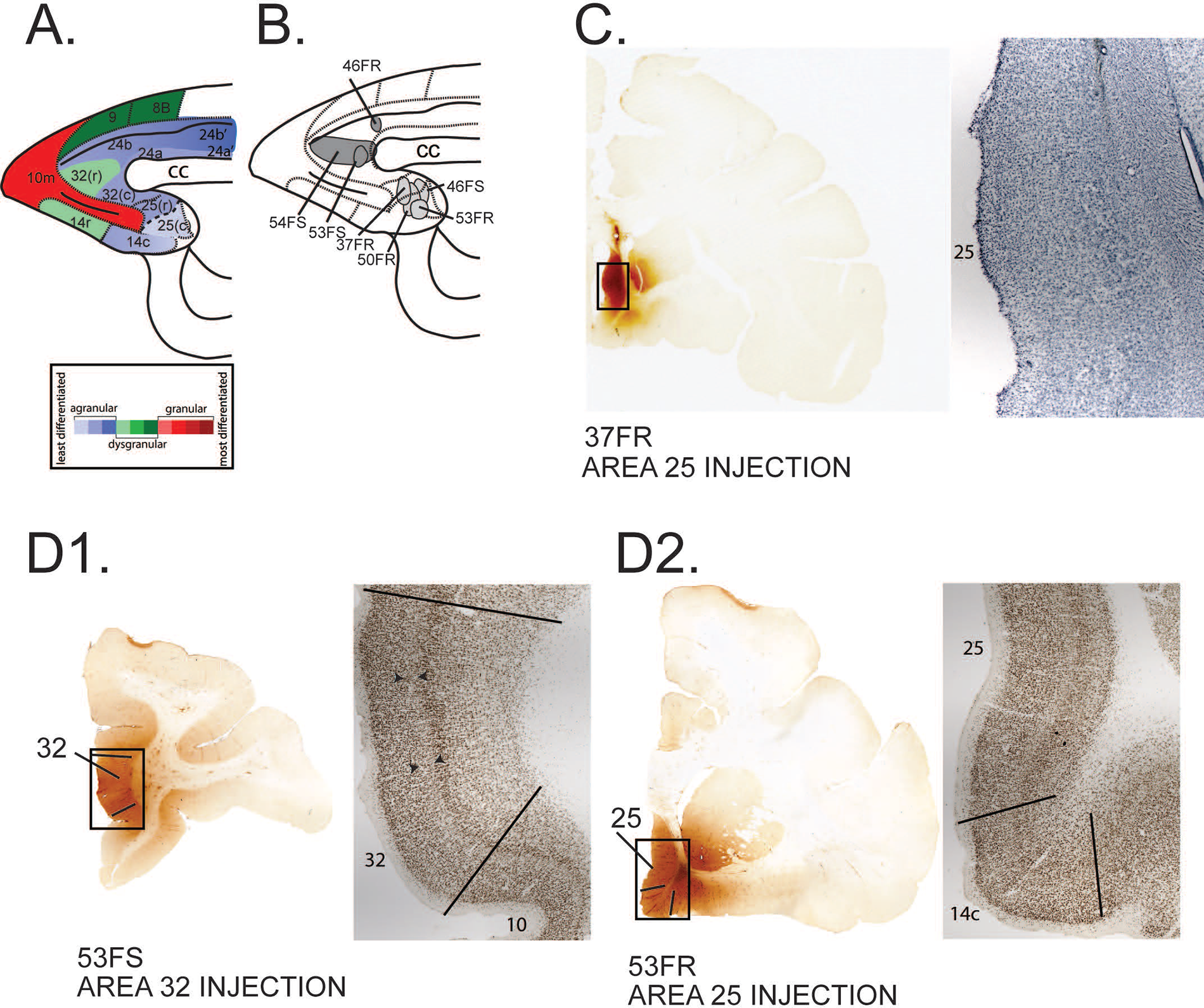
Locations of the sgACC (area 25/14c) and pgACC (area 24b, 32) injection sites in sagittal section of the prefrontal cortex. (assessed using cytoarchitectural criteria of Carmichael and Price, 1994). (A) Cytoarchitectural shifts across the medial prefrontal cortex, including the ACC. Agranular (blue), dysgranular (green) and granular (red). (B) Sagittal schematic of showing placement of anterograde injection sites. (C) Case 37FR, with a single injection into area 25r, and adjacent cresyl violet stained section. (D1, D2) Brightfield images of dual tracer injections into pgACC (area 32, 53FS) (D1) and sgACC (area 25/14c, 53FR) (D2) in the same hemisphere of Case 53. (Following the area 25 injection, anterogradely labeled fibers are seen leaving the cortex, and entering the ventral striatum). Adjacent NeuN-immunostained sections show respective the cytoarchitectural features at each site. Arrowheads delineate dysgranular layer IV in area 32 (D1).

### General topography of sgACC and pgACC inputs to the amygdala

All injections into the sgACC resulted in broad distribution of anterogradely labeled fibers, specifically in the medial nucleus (M), the lateral nucleus (L), the magnocellular subdivision of the accessory basal nucleus (ABmc), the magnocellular and intermediate subdivisions of the basal nucleus (Bmc, Bi), the medial subdivision of the central nucleus (CeM) and the amygdalostriatal areas (Fig. 4A-C, blue). Scattered labeled fibers existed in the parvicellular subdivision of the basal nucleus (Bpc) and accessory basal nucleus (ABpc), amygdalohippocampal area (AHA), and in the intercalated islands (Fig. 1E-F) that surrounded the basal nucleus, consistent with previous studies (Freedman et al., 2000). There were no apparent rostrocaudal differences in labeled fiber distribution, despite slight rostrocaudal differences in sgACC injection placement. Moreover, inclusion of the area 14c (in case 53FR) did not alter the distribution of labeled fibers in the amygdala.

**Figure 4.**
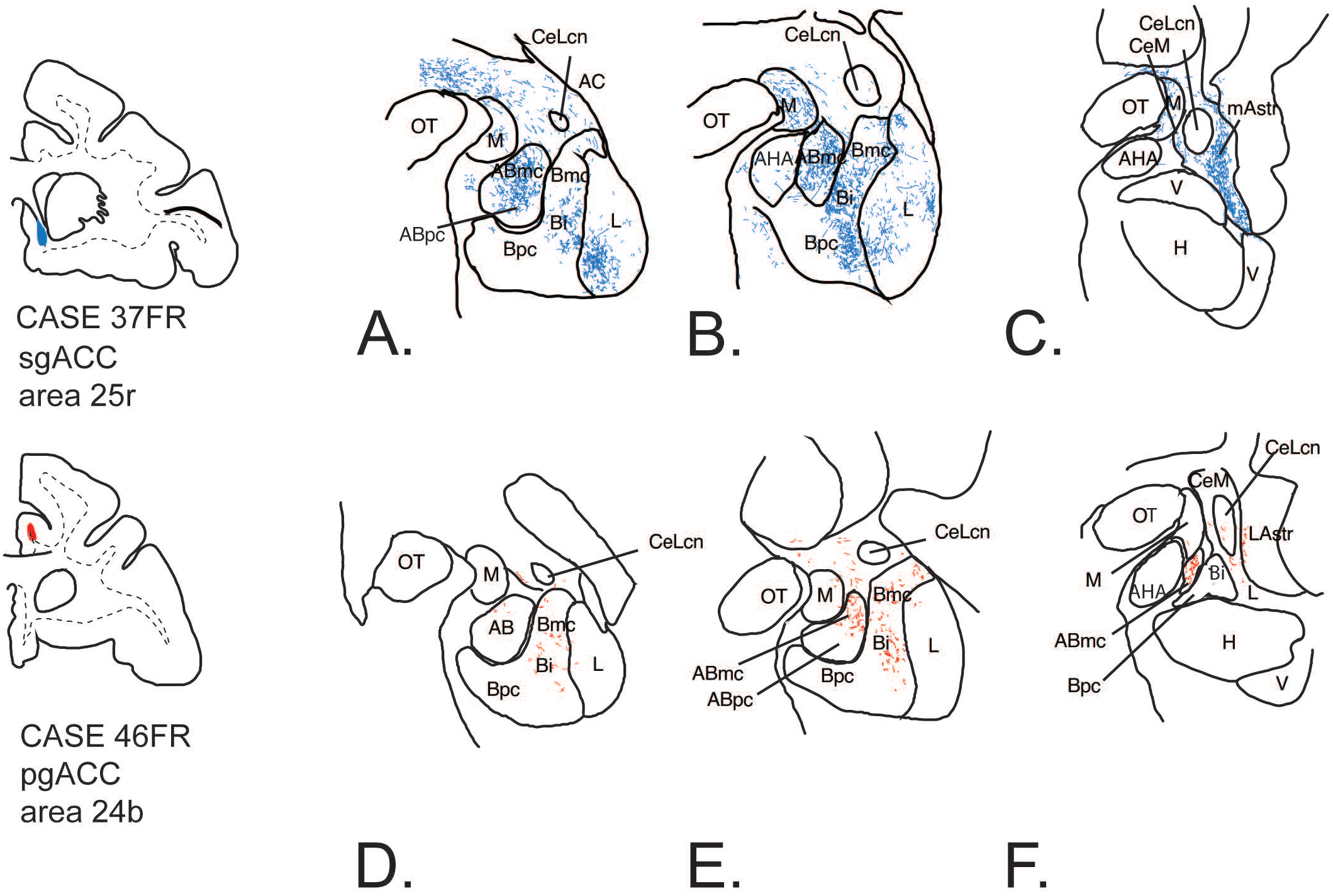
General topography of anterogradely labeled fibers in two different cases after injections into sgACC and pgACC. Fiber distribution following injections into sgACC (case 37FR, blue; A-C) and pgACC (case 46FR, red; D-F). ***Abbrs.* ABmc**, accessory basal nucleus, magnocellular subdivision; **ABpc**, accessory basal nucleus, parvicellular subdivision; **ABs**, accessory basal nucleus, sulcal subdivision; **AC**, anterior commissure; **AHA**, amygdalohippocampal region; **Bi**, basal nucleus, intermediate subdivision; **Bmc**, basal nucleus, magnocellular subdivision; **Bpc**, basal nucleus, parvicellular subdivision; **CeLcn**, lateral subdivision of the central nucleus; **H**, hippocampus; **L**, lateral nucleus; **M**, medial nucleus; **mAstr**, medial amygdalostriatal area; **OT**, optic tract; **P**, putamen; **V**, ventricle;

In general, tracer injections into the pgACC resulted in fewer labeled terminals than sgACC injections, and the distribution was more restricted (Fig. 4D-F, red). Because pgACC injections resulted in a more restricted pattern of anterogradely labeled terminals, compared with sgACC injections, we placed a single large injection encompassing area 32/24b (case 54FS) using three times the normal injection volume (150 nl) as a comparison. For this relatively large injection, labeled terminal fibers were still qualitatively less dense, and more confined, than sgACC injections using 40 nl (not shown). Labeled fibers were found in the accessory basal nucleus, magnocellular subdivision (ABmc) and the basal nucleus, intermediate and magnocellular subdivisions (Bi, Bmc) and in the medial and lateral subdivisions of the central nucleus (CeM, CeLcn,respectively), and amygdalostriatal area, with few to no labeled fibers found in other regions including the intercalated islands.

### pgACC and sgACC labeled fibers overlap in amygdala ‘hotspots’

Paired injections, matched for tracer volumes, were first mapped throughout the entire amygdala using adjacent sections containing labeled fibers from each ACC node (Cases 46 and Case 53, Fig. 5-1). The paired injection cases each had an injection into sgACC; case 46 had a companion injection in pgACC (area 24b), and case 53 had a companion injection in pgACC area 32. Tracers injected into the pgACC and sgACC had been ‘reversed’ in case 46 and 53 to control for possible effects of tracer transport (fluorescein injection into sgACC /fluororuby injections into pgACC for case 46; the reverse was done for case 53). Resulting maps were consistent with the patterns of labeled fibers seen in single-injection cases (Fig. 5-1), but permitted examination of patterns of segregation and convergence of labeled terminals from each site in the same animal.

Our analysis focused on all amygdala nuclei containing glutamatergic projection neurons (i.e. excluded the medial and central nucleus) (Fig. 5 A-F). As in single injection cases, sgACC labeled terminals targeted more nuclei (the accessory basal nucleus, basal nucleus, lateral nucleus, and AHA) compared to the pgACC (Fig. 5, A-C and D-F, blue). Labeled terminals from the pgACC injections targeted mainly the accessory basal nucleus and the basal nucleus, with no labeled fibers in the lateral nucleus or AHA (Fig. 5, A-C and D-F, red). Within the basal nucleus, the sgACC labeled fibers were found in all subdivisions, while pgACC labeled terminals were largely confined to the Bi (and to a lesser extent, the Bmc). In the accessory basal nucleus, sgACC labeled terminals extended into both the magnocellular and parvicellular subdivisions, whereas pgACC labeled fibers were confined to the magnocellular accessory basal nucleus. These results were consistent with previous retrograde data (Cho et al., 2013).

**Figure 5.**
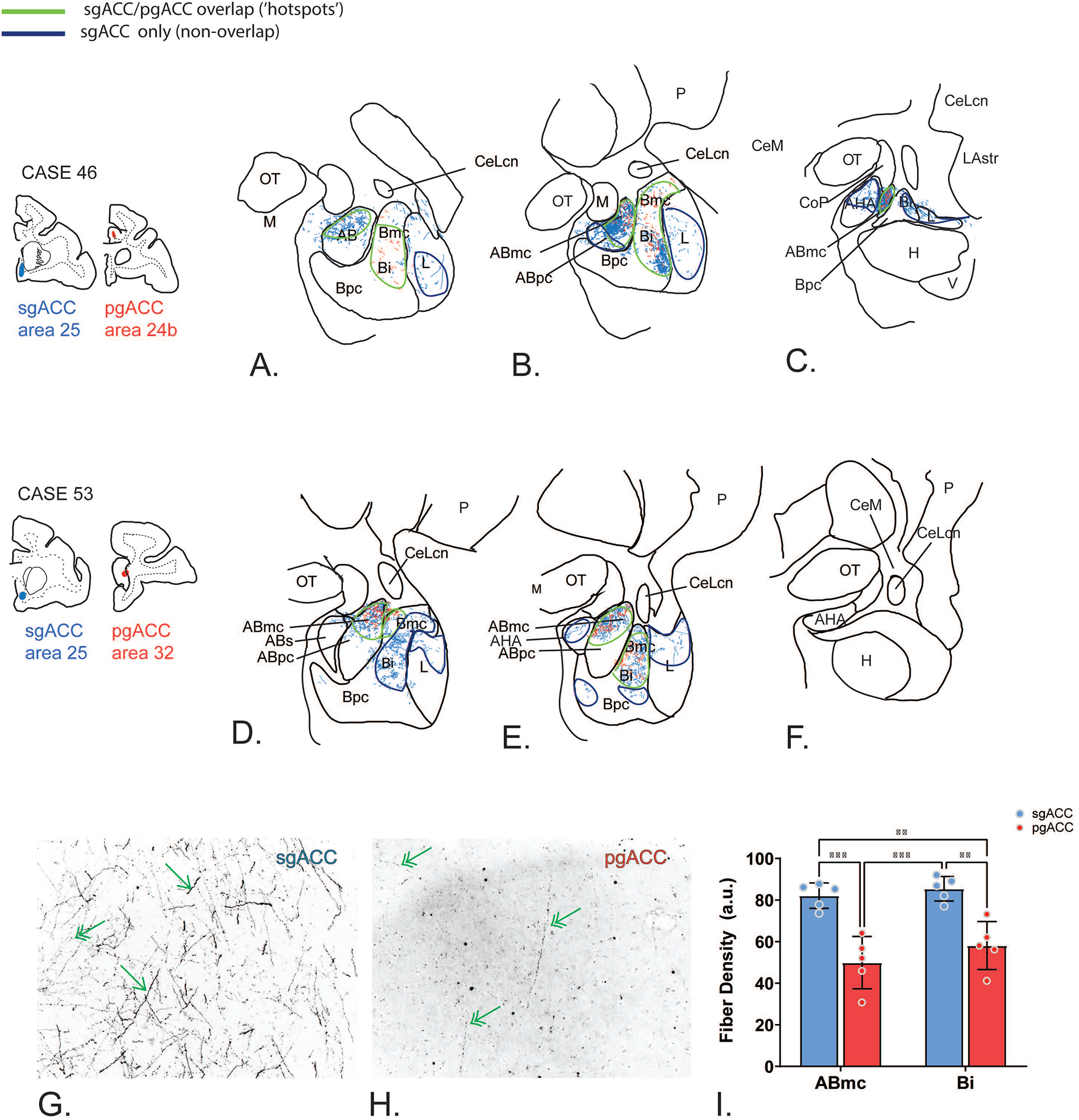
Anterogradely labeled fiber distribution in the main amygdala nuclei following paired pg- and sgACC tracer injections. Rostral to caudal distribution of sgACC (blue) and pgACC (red) fibers in Case 46 (A-C) and 53 (D-F). A-F. Areas of sgACc/pgACC overlap are circled in green; areas of sgACC non-overlap area circled in dark blue. G-H. Inverted dark-field micrograph of fiber termination in sgACC (G) and pgACC (H) at 20X magnification. Single headed arrows denote thick, beaded labeled fibers that were seen in the sgACC projection. Double-headed arrows show thin, beaded labeled fibers found in both the pgACC and sgACC projection field. Fiber density measures in each ‘hotspot’ for sgACC (blue) vs pgACC (red). See Extended data Figure 5.1 for labeled fiber distribution including the extended amygdala.

**Extended data Figure 5-1.**
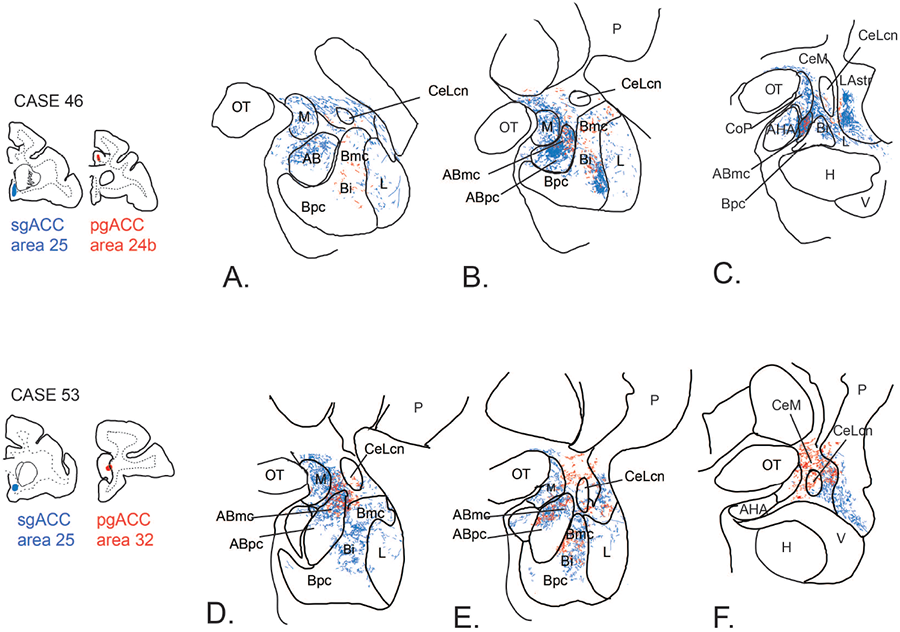
Anterogradely labeled fiber distribution following *paired* pg- and sgACC tracer injections, including extended amygdala regions. Rostral to caudal distribution of sgACC (blue) and pgACC (red) fibers in Case 46 (A-C) and 53 (D-F).

Overlap of pgACC and sgACC terminals occurred in areas where the pgACC had terminals: the intermediate subdivision of the basal nucleus and accessory basal nucleus (magnocellular subdivision) (Fig. 5A-C, and D-F, green outlines). That is, pgACC terminals were nested in the overall sgACC terminal distribution. We designated these regions of overlap ‘hotspots’. Regions where sgACC terminals did not overlap with pgACC labeled fibers were designated as segregated, or ‘non-overlapping’ regions, and also demarcated (dark blue outlines).

In ‘hotspots’, the relative density of labeled terminals following sgACC injections was generally higher than for pgACC injections, regardless of tracer used (Fig 5G-I). Some sgACC labeled beaded fibers were thicker, there were many thin, varicose labeled fibers (Fig. 5G). pgACC labeled fibers appeared thin and more diffuse (Fig. 5H). Quantification of fiber density confirmed significantly denser distributions of fibers originating from sgACC in both ABmc and Bi (‘hotspots’) compared to those originating from pgACC (Fig. 5I; sgACC:ABmc vs pgACC:ABmc= p=0.0003; sgACC:Bi vs pgACC:Bi= p=0.0018; Two-way ANOVA with Tukey’s multiple comparisons test). Further, sgACC-originating fiber density was significantly higher regardless of ‘hotspot’ region (sgACC:ABmc vs pgACC:Bi= p=0.0054; sgACC:Bi vs pgACC:ABmc=p=0.0001).

### Most CAMKIIα (+) cells in ‘hotspots’ receive dual contacts

For cases with paired injections (cases 46 and 53), we then conducted triple-labeling for each tracer and CAMKIIα (a marker of pyramidal neurons in the amygdala (McDonald et al., 2002)). We used confocal methods to determine the degree of fiber ‘contact’ onto putative excitatory neurons in both ABmc and Bi ‘hotspots’ where labeled terminals from both the sgACC and pgACC converged (Fig. 6A-B; see Methods). Thresholding of randomly selected CAMKIIα(+) cells and tracer labeled fibers was completed in separate channels, and overlaid.

**Figure 6.**
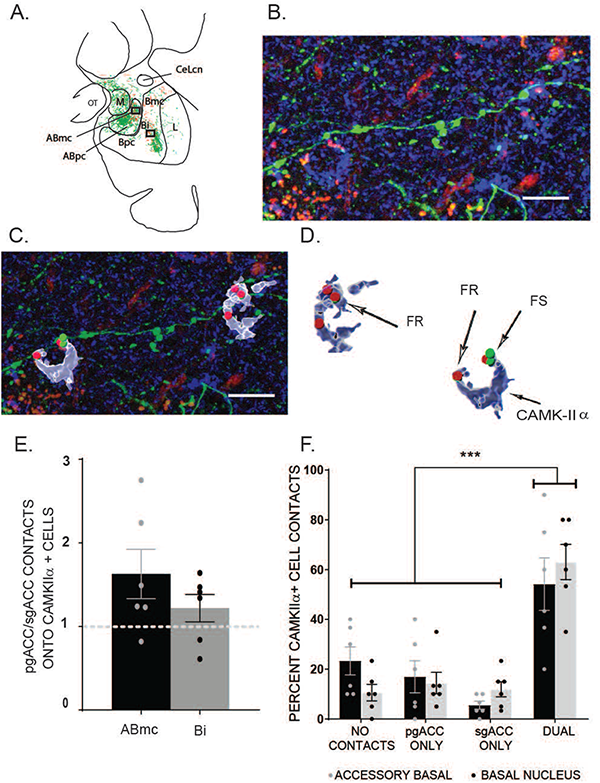
Triple immunofluorescent analysis and quantification of tracer contacts following paired pgACC/sgACC injections. (A) Camera lucida rendering of anterograde fiber distribution following pgACC (red) and sgACC (green) injections, used for selection and registration of ‘hotspots’ in the ABmc and Bi (boxed areas). (B) CAMK-IIα (blue), fluoroscein tracer (FS) and fluoruby tracer (FR) triple immunofluorescent confocal micrograph collected in the ABmc ‘hotpspot’, boxed area in A. Scale bar = 10 um (C) Overlay of Imaris software ‘surface’ (CAMK-IIα) and ‘spot’ (tracer) renderings onto confocal micrograph (shown in B). Scale bar = 10 um (D) Imaris rendered objects depicting sites of tracer contact with CAMK-IIα cells. (E) The proportion of pgACC/sgACC contacts onto all CAMK-IIα cells counted in the ABmc and Bi ‘hotspots’. (F) The proportion of CAMK-IIα cells with no contact, single contacts, or dual contacts from pgACC and sgACC anterogradely labeled fibers in the ABmc and Bi (****= p<0.001, two-way ANOVA with Tukey multiple comparisons post hoc test). See Extended data Figure 6-1 for raw counts and ratios of sgACC and pgACC contacts by region of interest and rostrocaudal level, and Extended data Figure 6-2 for pooled raw counts and ratios of pgACC and sgACC contacts by case and region of interest.

**Extended data Figure 6-1.**
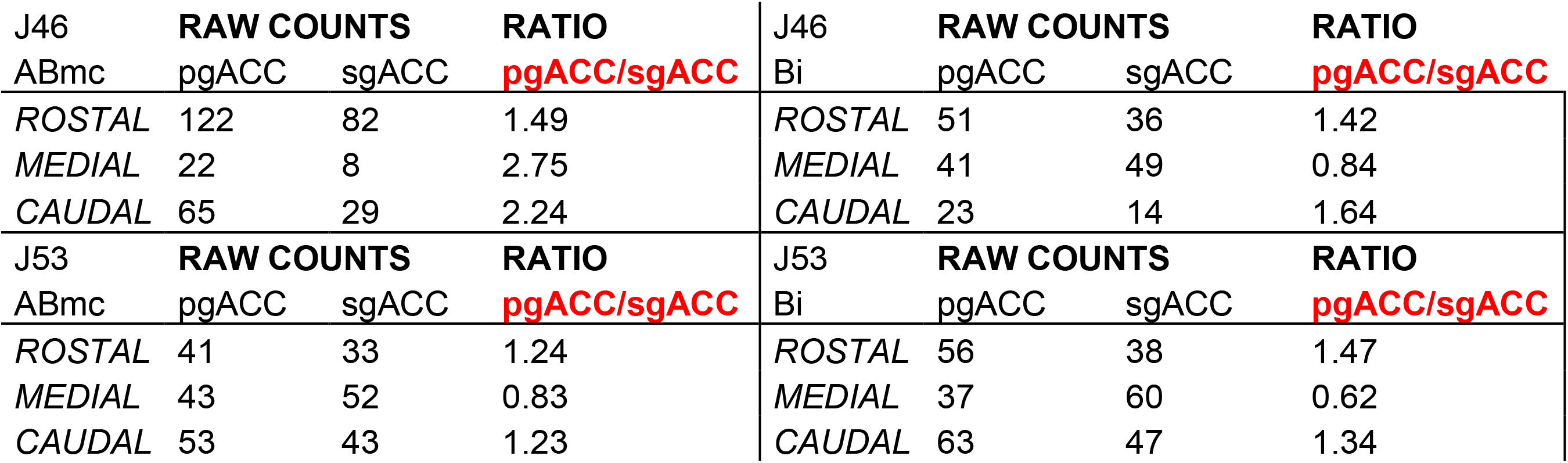
Rostro-caudal levels: Raw counts and ratios of sgACC and pgACC contacts onto CAMK-IIα cells in each case by region of interest at 3 rostrocaudal levels. No rostrocaudal differences were found. Two-way ANOVA with Tukey Multiple Comparisons test, F(5,16)= 0.2176, p=0.9495).

**Extended data Figure 6-2.**
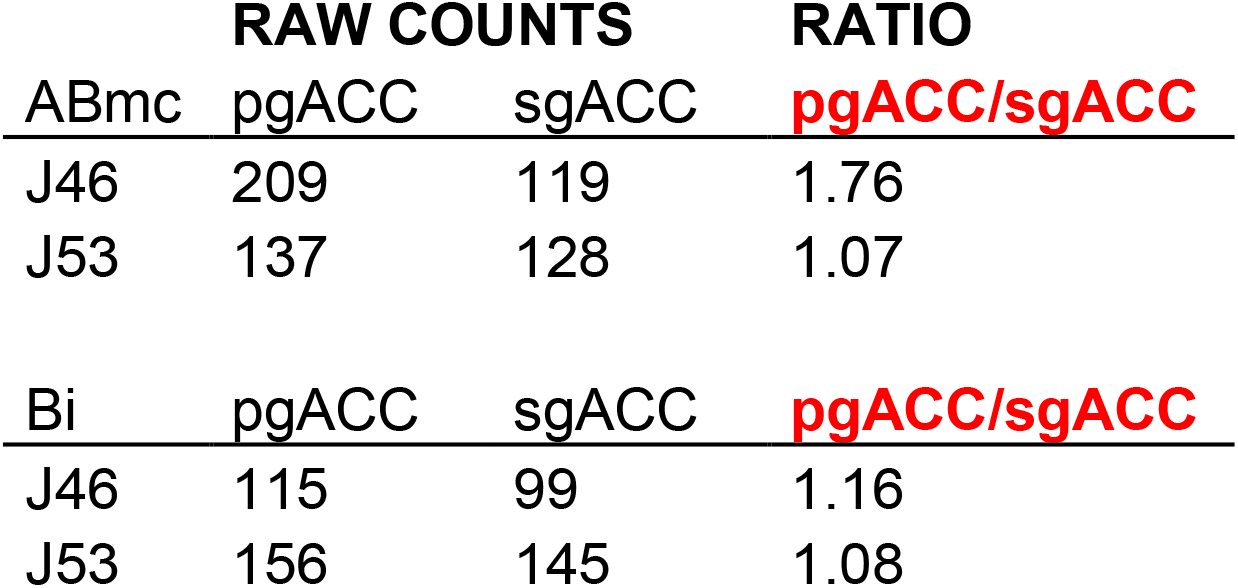
Raw counts and ratios of pgACC and sgACC contacts onto CAMK-IIα cells by case and region of interest. No rostrocaudal differences were found. One-Way ANOVA with Tukey multiple comparisons tests, Case 46:ABmc= F(2,4)=11.85, p=0.0609; Case 46:Bi= F(2,3)=6.231, p=0.0855; Case 53:ABmc= F(2,5)=5.136, p=0.0613, p=0.0613; Case 53:Bi= F(2,6)=0.5588, p=0.5990)

Randomly selected CAMK-IIα(+) cells in each ‘hotspot’ in each case were examined for contacts with either sgACC or pgACC tracer (+) boutons (Fig. 6C-D). Given some variability in labeled fiber distribution across the rostrocaudal extent of the ABmc and Bi for all tracers/injections at the macroscopic level, we first examined whether there were differences in numbers of pgACC and sgACC contacts across the entire rostrocaudal expanse in each animal. No rostrocaudal differences were found (see Fig. 6-1 for rostral-caudal raw data; Two-way ANOVA with Tukey Multiple Comparisons test, F(5,16)= 0.2176, p=0.9495), and results of all sections were grouped for each animal (Fig. 6-2, pooled raw data). For all CAMK-IIα(+) cells examined (n=300), there were no significant differences in the number of pgACC and sgACC contacts (sgACC: 247 contacts in ABmc, 244 contacts in Bi; pgACC: 346 contacts in ABmc, 271 contacts in Bi; ANOVA with Tukey’s Multiple Comparisons test; F(3,4)=0.9985, p=0.4795; n.s.). The ratio of pgACC to sgACC labeled contacts was approximately equal in both the ABmc and Bi ‘hotspots (Fig.6E, n= 150 total cells ABmc, n=150 total cells Bi, two-tailed students t-test; p=0.2525; n.s.).

We then explored the extent to which pgACC and sgACC contacts converged on the same CAMK-IIα(+) cells in each area. The majority of randomly selected CAMK-IIα(+) neurons had contacts from both the sgACC and pgACC, in both ABmc and Bi regions (Fig. 6F, Two-way ANOVA with Tukey multiple comparisons test; F(7,24)= 9.634; DUAL vs all other contact profiles= ***= p<0.0001). Relatively lower proportions of CAMK-IIα(+) neurons had no contacts from either projection or had contacts from a single projection. There were no significant differences in the distribution of dual-contact CAMK-IIα(+) neurons in the rostrocaudal plane (One-Way ANOVA with Tukey multiple comparisons tests, Case 46:ABmc= F(2,4)=11.85, p=0.0609; Case 46:Bi= F(2,3)=6.231, p=0.0855; Case 53:ABmc= F(2,5)=5.136, p=0.0613, p=0.0613; Case 53:Bi= F(2,6)=0.5588, p=0.5990).

### The proportion of pgACC to sgACC dual contacts are tightly balanced

Since the majority of CAMK-IIα(+) neurons in ‘hotspots’ had dual contacts, we calculated the relative weighting of sgACC and pgACC contacts on individual dually contacted cells, and the distribution of the ratios of pgACC-to-sgACC contacts throughout the ABmc and Bi populations (Fig. 7 A-B, Fig. 7-1). Ratios of contacts from each area onto individual CAMK-IIα(+) neurons ranged from 0 to 3.5. The majority of dual pgACC: sgACC contact ratios for both the ABmc and Bi were in a narrow 1.0-1.5 range, suggesting a relatively tight balance of inputs onto individual pyramidal neurons in most cases. Given this narrow range, we pooled our data and performed quantification based on ratio bin counts (Fig. 7C). Quantitative comparisons of ratio bin counts display significantly higher counts in the range of 1-1.5 when compared to every other bin (*=p<0.05, ****= p<0.001, one-way ANOVA with Tukey multiple comparisons post hoc test).

**Figure 7.**
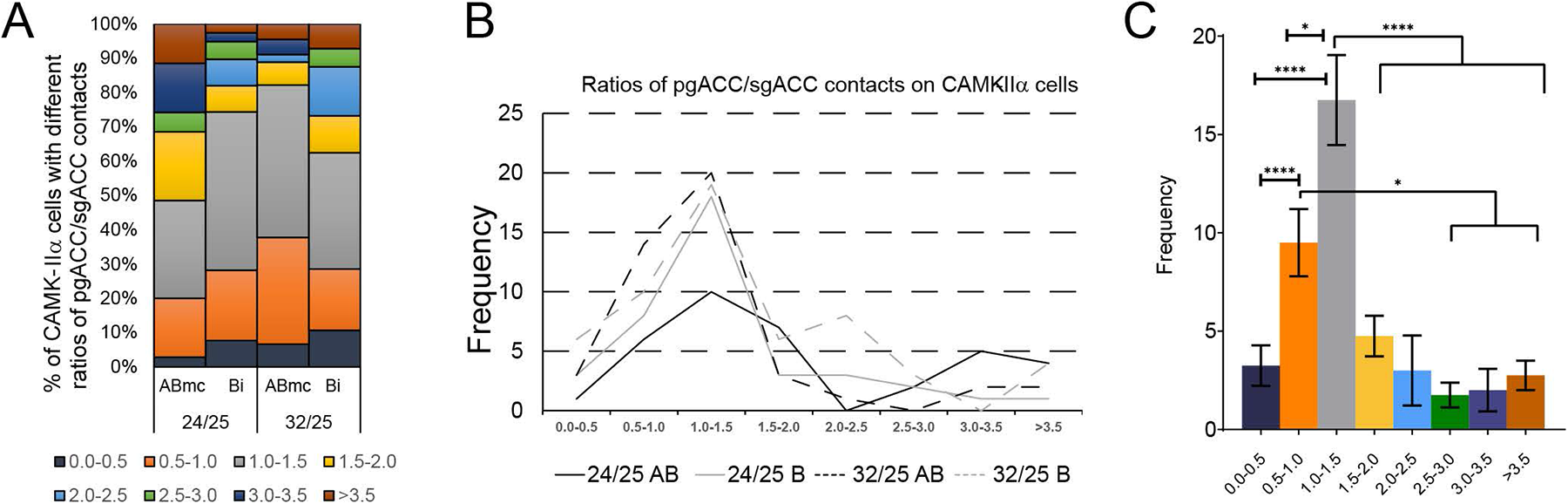
The proportion of pgACC to sgACC dual contacts are tightly balanced in amygdalar hotspots. (A) Binned ratios of pgACC/sgACC contacts onto individual dual-contacted CAMK-IIα cells. A large percentage of CAMK-IIα cells had ratios 1:1 pgACC-to-sgACC contacts (gray bars) in both the ABmc and Bi. (B) Frequency distribution of pgACC/sgACC contacts onto CAMK-IIα cells. Ratio bins range from 0 to >3.5. (C) Quantitative comparisons of ratio bin counts (*=p<0.05,****= p<0.001, one-way ANOVA with Tukey multiple comparisons post hoc test). See Extended data Figure 7-1 for raw bin counts of dual contact per CAMK-IIα cell by case and region, and quantitative comparisons across bins.

**Extended data Figure 7-1.**
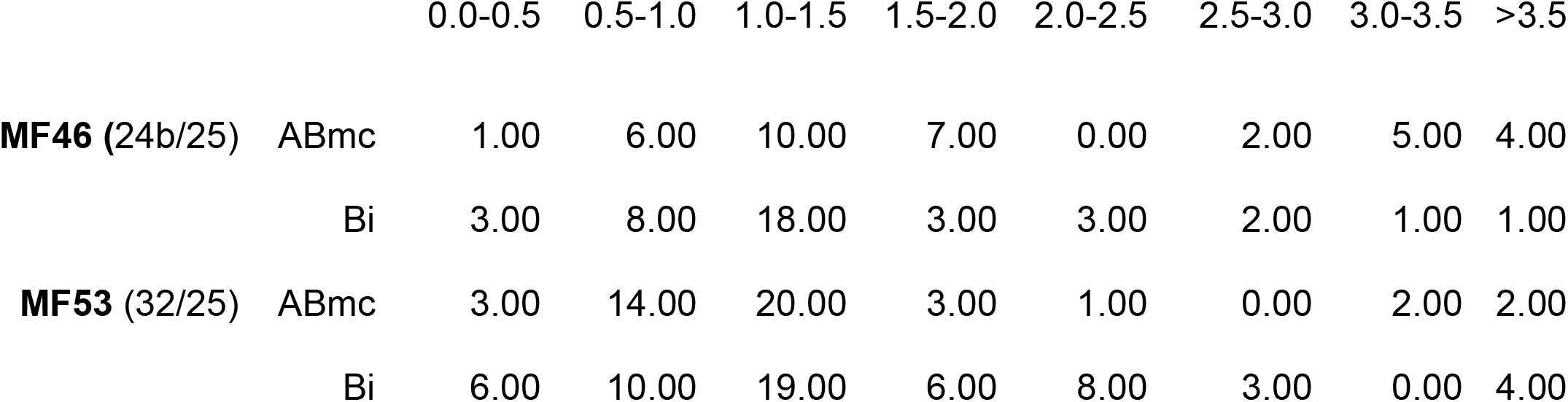
Raw bin counts of pgACC/sgACC dual contacts onto CAMK-IIα cells by case and region of interest. Quantitative comparisons of ratio bin counts display significantly higher counts in the range of 1-1.5 when compared to every other bin (*=p<0.05, ****= p<0.001, one-way ANOVA with Tukey multiple comparisons post hoc test).

### sgACC and pgACC synaptic bouton volumes

We next examined bouton volumes from the sgACC and pgACC in each ‘hotspot’ as an approximation of synaptic ‘strength’ (Petrof and Sherman, 2013). sgACC non-overlapping (non-OL) terminals in the lateral nucleus and AHA were examined for comparison. Data were collected on terminal boutons and bouton*s ‘en passant’* using stereologic methods in adjacent, single labeled sections. In all areas assessed, the majority of boutons were significantly less than 0.52 um^3^ volume (equivalent 1um diameter) for sgACC and pgACC afferents (Fig. 8 A-E; multiple comparison tests were performed to compare small vs. large bouton across region and case. In all situations, there were significantly more small boutons compared to large boutons (p=0.008 [J46], p=0.015 [J53], p=0.005 [pooled])). However, in ‘hotspots’ in the ABmc and Bi, there was a higher frequency of relatively large terminals for the sgACC (Fig. 8-1 and 8-2).

**Figure 8.**
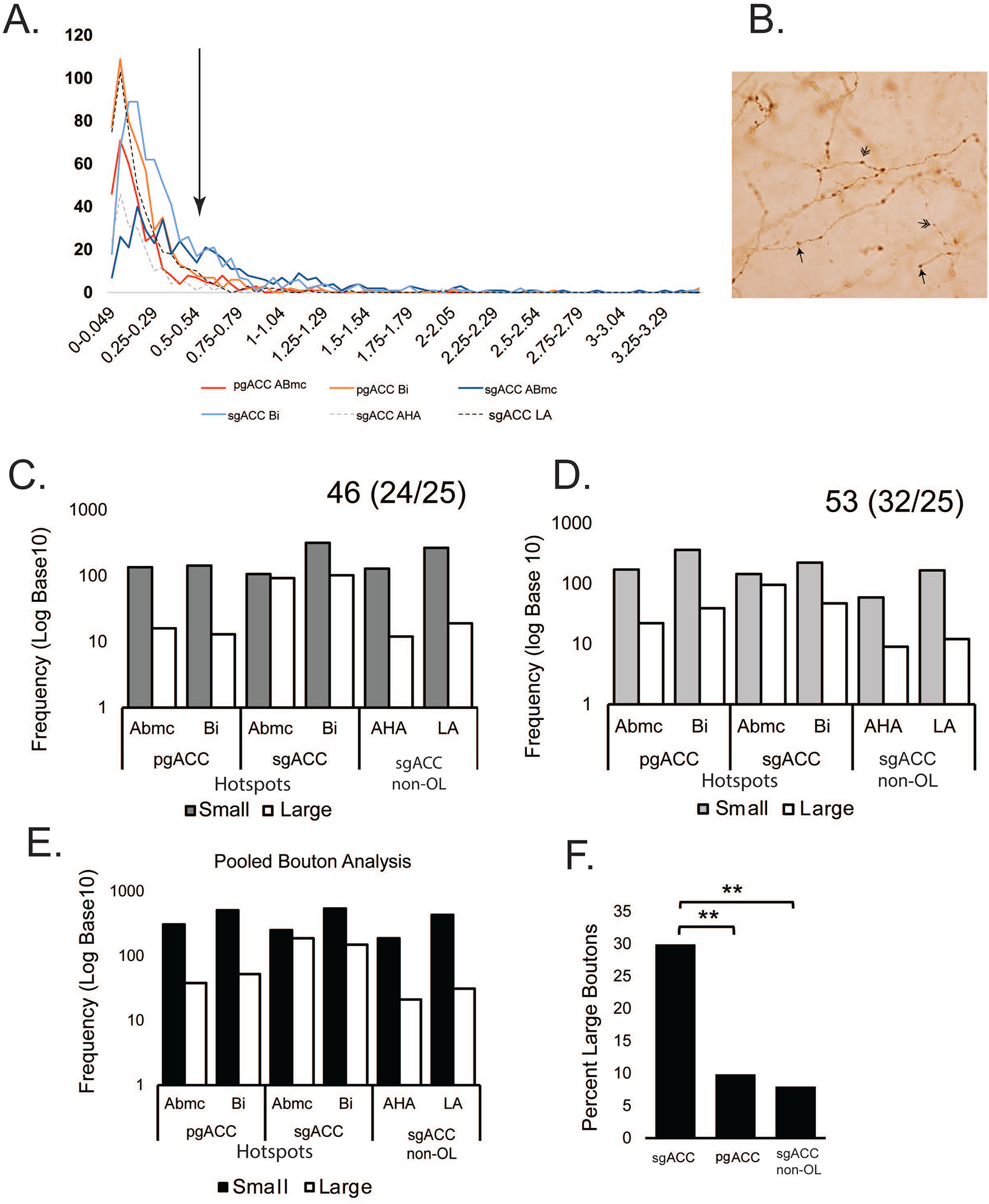
Unbiased stereology and quantitative analysis of anterogradely labeled synaptic boutons at 100x. (A) Frequency bin representation of volumetric measurements of tracer-labeled synaptic boutons from the sgACC (blue) and pgACC (red) in ‘hotspots’, and sgACC terminal boutons in non-overlapping AHA (gray dashed) and Lateral nucleus regions (black dashed) (for comparison). Volume bins ranged from 0->3.45 um^3^. Many boutons fell below 0.52 um^3^ for all projections (arrow denotes shift to large categorized boutons). (B) A representative brightfield micrograph of anterograde tracer-labeled fibers in the basal nucleus ‘hotspot’ showing bouton types. Terminal boutons (single arrowhead) were characterized by a distinct swelling with an apparent stalk emanating from the axon terminal. *En passant boutons* (double arrowhead) displayed a characteristic swelling along the terminal fiber. Scale bar= 25 um. (C-D) Log frequency of small (black bars) vs large bouton (white bars) comparisons for each case with paired injections: (C) Case 46 (areas 24/25) (D) Case 53 (areas 32/25) (E) Pooled analyses, both cases. Multiple comparison tests were performed to compare small vs. large bouton across region and case. In all situations, there were significantly more small boutons compared to large boutons (p=0.008 [J46], p=0.015 [J53], p=0.005 [pooled]). Large boutons (greater than 0.52 um^3^ volume) were significantly more frequent on axons originating from sgACC in hotspots, but not non-overlapping regions. (One-way ANOVA with Tukey multiple comparisons, F(2,3)= 96.18, p= 0.0019; sgACC vs pgACC= p=0.0030; sgACC vs non-OL= p=0.0023). Extended data Figure 8-1 shows raw data counts of bouton type. Extended data Figure 8-2 shows regional counts of bouton size across injection source (pgACC, 32/24; sgACC, 25; sgACC, non-OL regions) and ROI (ABmc, Bi, AHA and LA).

**Extended data Figure 8-1.**
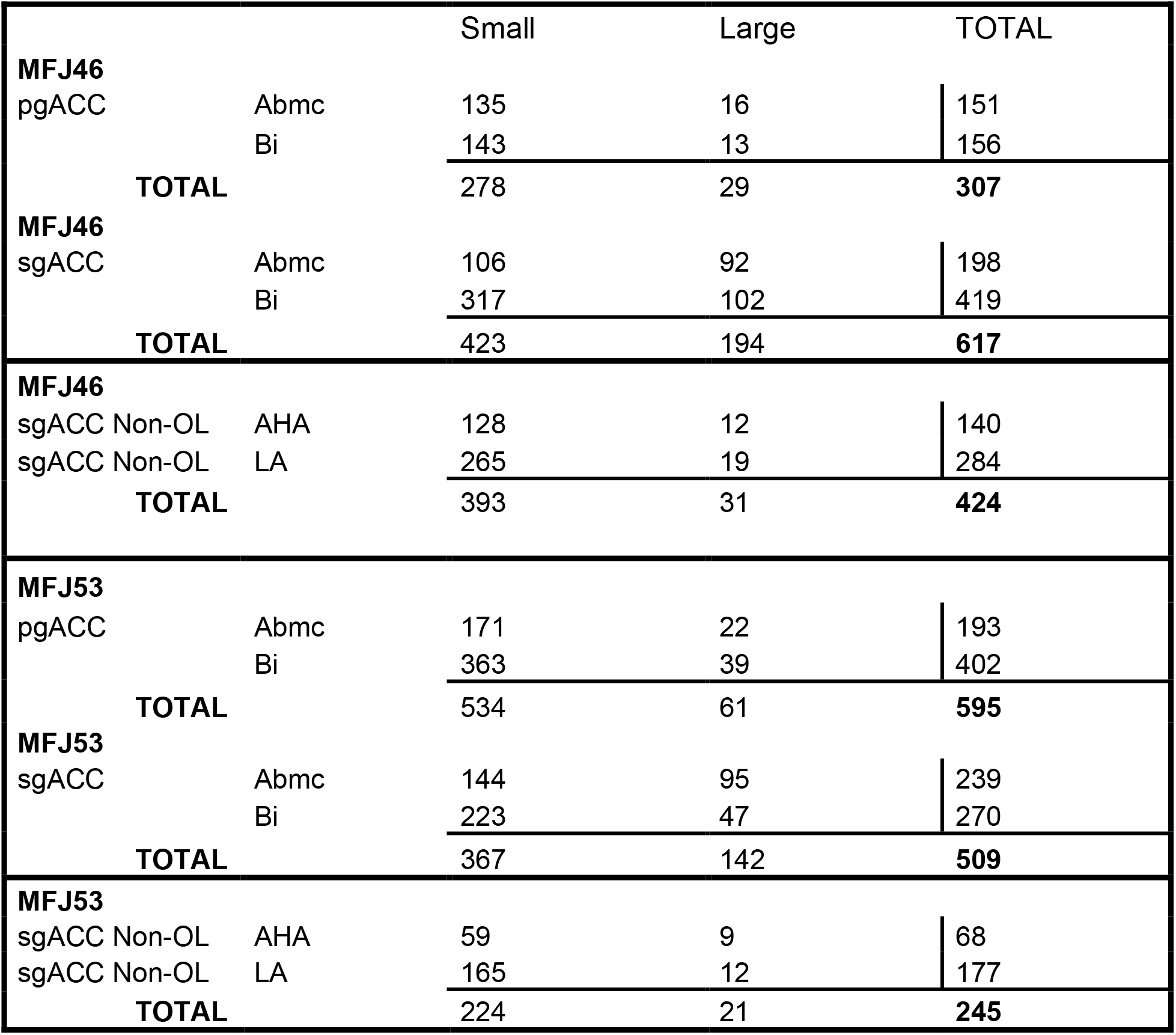
Regional counts of small (<0.52 um^3^) and large (>0.52 um^3^) boutons across paired cases (Case 46, 24/25 injections; Case 53, 32/25 injections), injection source (pgACC, 32/24; sqACC, 25; sgACC (non-OL; AHA and LA) and ROI (ABmc, Bi, AHA, and LA). We found that the sgACC terminals that overlapped pgACC-containing sites had significantly more large boutons (27-31%) compared to the pgACC (9-10%) [*sgACC*: J46 (total large boutons/total) = 194/617; M53 (total large boutons/total) = 142/509; *pgACC*: J46 (total large boutons/total) = 29/307; M53 (total large boutons/total) = 61/595; p=0.003; One-Way ANOVA with Tukey’s multiple comparison test, F(2,3)=96.18= p=0.0019; sgACC vs pgACC= p=0.0003].

**Extended data Figure 8-2.**
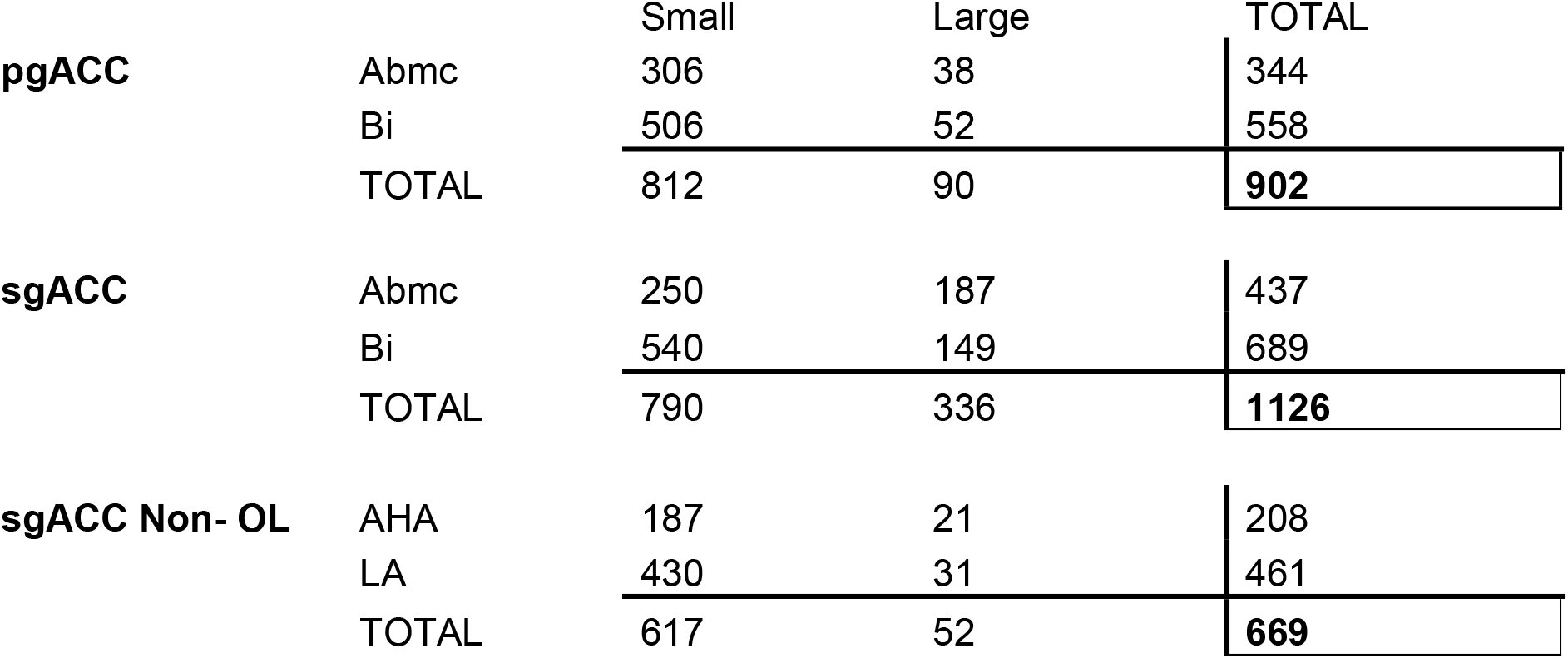
Regional counts of small (<0.52 um^3^) and large (>0.52 um^3^) boutons across injection source (pgACC, 32/24; sgACC, 25; sgACC, non-OL regions) and ROI (ABmc, Bi, AHA and LA). ‘Large’ bouton comparisons from the same afferent source did not differ significantly across the ABmc, Bi (p=0.0.3739, Two-way ANOVA, F(2,2)=1.674). sgACC non-OL areas had significantly fewer large boutons (Tukey Multiple Comparisons test following one-way ANOVA; sgACC vs non-OL= p=0.0023).

Using >0.52 um^3^ (> 1 um diameter) as a cut-off for ‘large’ boutons, we found that the sgACC terminals that overlapped pgACC-containing sites had significantly more large boutons (27-31%) compared to the pgACC (9-10%) (Fig. 8F; *sgACC*: J46 (total large boutons/total) = 194/617; M53 (total large boutons/total) = 142/509; *pgACC*: J46 (total large boutons/total) = 29/307; M53 (total large boutons/total) = 61/595; p=0.003; One-Way ANOVA with Tukey’s multiple comparison test, F(2,3)=96.18= p=0.0019; sgACC vs pgACC= p=0.0003). ‘Large’ bouton comparisons from the same afferent source did not differ significantly across the ABmc, Bi (p=0.0.3739, Two-way ANOVA, F(2,2)=1.674), suggesting a common feature of sgACC inputs in ‘hotspots’. In contrast, sgACC non-OL areas had significantly fewer large boutons, suggesting a specific feature in ‘hotspots’ (Fig 8F, Tukey Multiple Comparisons test following one-way ANOVA; sgACC vs non-OL= p=0.0023).

## Discussion

In broad-based studies, we previously showed that prefrontal-amygdala paths are organized in hierarchical arrays, dictated by the degree of laminar differentiation of the cortex (Cho et al., 2013). To examine this relationship in a more focused way and at the cellular level, we placed anterograde tracer injections into two different nodes of the ACC that have progressive laminar features (pgACC>sgACC), and distinct connections and functions (Carmichael and Price, 1995; Rushworth et al., 2013; Neubert et al., 2015; Palomero-Gallagher et al., 2015). In this study, we found that at the ‘meso-scale’ that pgACC afferent terminals are always ‘nested’ in broader sgACC terminals in the basal and accessory basal nuclei, confirming previous retrograde results, and elucidating connectional principles of the two ACC-amygdala microcircuits.

At the cellular level, we found that the majority of CAMIIα (+) amygdala neurons (putative projection neurons) in ‘hotspots’ of convergence were co-contacted by terminals from the sgACC and pgACC. This was true regardless of whether the ‘hotspot’ was in the ABmc or Bi. Despite the size of the ‘hotspots’ in the large primate samples, there were no rostrocaudal differences for these findings. Another key finding is that the ratio of pgACC-to-sgACC contacts was highly consistent within and across ‘hotspots’ and fell mainly in the range of 1.0-1.5. This suggests a general consistency in the relationship in pgACC:sgACC afferent balance onto common post-synaptic cells in regions of convergence, at least in young Macaques. Finally, sgACC terminals in ‘hotspots’ were more likely to have large boutons, compared to pgACC terminals, in the zones of convergence, suggesting possible differences in transmission speed and efficiency.

### Layering of amygdala subcircuits

The agranular sgACC is strongly interconnected with the midline thalamus, hypothalamus and periacqueductal gray, all of which mediate arousing and autonomic components of emotional responses (Van Hoesen et al., 1993; Vogt, 2005; Rudebeck et al., 2014). The sgACC has therefore been considered a core node in the ‘somatic’ marker hypothesis, which states that covert signals from the body (such as autononomic features and visceral sensations) are important in shaping emotions and, eventually, action (Damasio, 1996; Chudasama et al., 2013). In contrast, the pgACC (Area 24/32) sits directly and dorsally adjacent to the sgACC and is more organized in its laminar construction. However, the pgACC does not have direct connections to the internal milieu via midline connections like the sgACC, although it shares many of the same ‘limbic’ connections and receives input from the sgACC (Barbas et al., 1999; Joyce and Barbas, 2018; Sharma et al., 2020). The pgACC plays a prominent role in rapid assessment of competing choices (including ‘conflictual’ choices) and decision-making, based on studies in monkeys (Rudebeck et al., 2006; Amemori and Graybiel, 2012; Pryluk et al., 2020) and in the human (Etkin et al., 2006; Modirrousta and Fellows, 2008; Maier and di Pellegrino, 2012; Ito et al., 2017). Not surprisingly, the pgACC is activated during tasks involving social decision-making tasks, which are intuitive and rapid and involve predicting outcomes based on social cues such as facial expression (Apps et al., 2016; Dal Monte et al., 2020). The fact that ‘salience’-detecting (sgACC) and ‘social/conflict-monitoring’ (pgACC) components of the ACC have an overlapping, afferent influence in specific ‘hotspots’ of the ABmc and Bi, suggests that the internal ‘salience’ information co-regulates amygdala neurons involved in decision-making networks.

The amygdala is involved in an array of functions including fear conditioning and extinction (Phelps et al., 2004; Quirk and Beer, 2006); safety signaling (Genud-Gabai et al., 2013); updating value representations (Buchel et al., 1998); responses to emotion in facial expressions (Breiter et al., 1996; Morris et al., 1996; Fitzgerald et al., 2006), and social decision-making and behavior (Chang et al., 2015; Minxha et al., 2017; Gothard et al., 2018). Current evidence from chronic recordings in monkeys indicates that the amygdala’s capacity to participate in all these tasks and contexts is due to a capacity for multi-dimensional processing (Putnam and Gothard, 2019; Pryluk et al., 2020). For example, the same amygdala populations that respond to direct eye gaze (a threat cue in nonhuman primates; Kalin and Shelton, 1989) also respond to non-social aversive stimuli (air-puff). Conversely, neurons that respond to averted eye gaze (a cue predicting submissive, positive social interactions) also respond to juice reward. Amygdala neurons are thus able to flexibly code across social and nonsocial stimuli to predict outcome in specific contexts. The present anatomic findings suggest that this multi-dimensionality may be served by the high rate of converging contacts on the same amygdala neurons from functionally distinct ACC regions, which may confer flexibility across various stimuli and contexts.

### High convergence of sgACC/pgACC contacts on projection neurons

The majority (85%) of cortical afferent inputs terminate onto pyramidal neurons, based on work in rodents (Brinley-Reed et al., 1995). Synapses located closest to the soma (or on the soma itself) have the most influence on the overall circuit, as they act to strengthen or weaken the overall cellular response (reviewed in; Villa and Nedivi, 2016). One of our most striking findings was that sgACC and pgACC terminals target the same pyramidal cell populations, with relatively few pyramidal neurons having only a single contact. Consistent with this finding, functionally related excitatory inputs are often clustered in regards to synaptic placement along the cell and are thought to possibly contribute to coordinated regulation of synaptic plasticity among co-active inputs (DeBello et al., 2014). Through spatial clustering, temporally co-activated excitatory inputs are more likely to initiate a potentiating response within the cell than inputs spaced a distance from one another. Although we did not examine synaptic contacts onto distal dendrites, the finding that dual contacts at the soma/proximal dendrites were the rule, and were tightly balanced in relationship to one another, strongly suggests cooperative actions in regulating post-synaptic cell excitability. The pgACC and sgACC are separate, but closely related brain regions with respect to connectivity and function. They therefore follow this general principle of “coordinated regulation” by functionally related inputs, contributing to the integration of complex or fluid informational streams.

The highly consistent ratio of sgACC to pgACC contacts in normal young animals raises questions about how and when this precise ratio is developed. In mice, the long-range afferents from the medial prefrontal cortex in general arrive in the basal nucleus by postnatal day (PND 10-15) (Bouwmeester et al., 2002; Arruda-Carvalho et al., 2017), but synaptic strength continues to develop until PND 30 (early adolescence) based on physiologic data. While the development of the relative ‘balance’ of the sgACC and pgACC (infralimbic and prelimbic cortices in rodents) terminals on the post-synaptic neuron is not known, experience-dependent plasticity likely contributes (Holtmaat et al., 2006). We speculate that experiences in early life contribute to sgACC:pgACC terminal balance.

A factor in the function of terminals is synapse size. Glutamatergic synapses are classified broadly into Class I and Class II synapses (Petrof and Sherman, 2013). Class I synapses are large (> 1 um diameter), associated with large axons, and release glutamate to affect ‘all or none’ action potentials. They are considered ‘drivers’ of the circuit. Larger glutamatergic synapses are associated with larger post-synaptic densities (Sheng and Kim, 2011), larger axon diameters (Innocenti and Caminiti, 2017) and greater transmitter release (Rosenmund and Stevens, 1996; Murthy et al., 1997). These structural specializations are thought to prioritize functional inputs in terms of timing and signal strength. In contrast, Class II synapses are smaller and modulatory, and shape post-synaptic excitability. In the present study, a higher number of large boutons in sgACC terminals in ABmc and Bi ‘hotspots’ may confer ‘driver’ function, balanced by pgACC modulators. In future studies, a key question will be to describe sgACC versus pgACC inputs at the ultrastructural level; to determine synaptic size, location (i.e. axo-dendritic spine/shaft, axo-somatic, etc.), and number (i.e single or multiple bouton contacts) (reviewed in, Yang et al., 2018) associated with glutamatergic post-synaptic neurons (Cover and Mathur, 2020).

## Conclusion

The primate amygdala is evolutionarily expanded, accompanied by increased levels of cellular complexity and coding capacity (Grabenhorst et al., 2019; Dal Monte et al., 2020; Gothard, 2020). Consistent with its cellular complexity, divergent coding schemes in the primate amygdala operate to facilitate different functions (Pryluk et al., 2020). Our results may help explain multi-dimensional coding flexibility of amygdala neurons since broad-based sgACC terminal fields appear capable of driving wide-spread activity across the main amygdala nuclei. In contrast, pgACC inputs follow a more restricted, nested topography (Cho et al., 2013) (present results), forming highly convergent contacts on sgACC-recipient neurons in specific subregions. This arrangement allows maximum flexibility, with the sgACC providing broad ‘internal salience’ feedback to the amygdala, but also permitting cooperative signalling with pgACC inputs for conflict monitoring and social decision-making in ‘hotspots’ of convergence.

## Data sharing statement

Digital data can be accessed by contacting the corresponding author at.

## Acknowledgments

This work was funded through the support of the National Institute of Mental Health R01MH63291 (J.L.F.). We thank Nanette Alcock for assistance with histology and immunohistochemistry.

